# Pneumococcal genetic variability influences age-dependent bacterial carriage

**DOI:** 10.1101/2021.03.03.433546

**Authors:** PHC Kremer, B Ferwerda, HJ Bootsma, NY Rots, AJ Wijmega-Monsuur, EAM Sanders, K Trzciński, AL Wyllie, P Turner, A van der Ende, MC Brouwer, SD Bentley, D van de Beek, JA Lees

## Abstract

The pneumococcal conjugate vaccine (PCV) primarily reduces disease burden in adults through a reduction in carriage prevalence of invasive serotypes in children. Current vaccine formulations are the same for both adults and children, but tailoring these formulations to age category could optimize vaccine efficacy. Identification of specific pneumococcal genetic factors associated with carriage in younger or older age groups may suggest alternative formulations and contribute to a better mechanistic understanding of immunity. Here, we used whole genome sequencing to dissect pneumococcal variation associated with age. We performed genome sequencing in a large carriage cohort, and conducted a meta-analysis with an existing carriage study. We compiled a dictionary of pathogen genetic variation including serotype, sequence cluster, sequence elements, SNPs, burden combined rare variants, and clusters of orthologous genes (COGs) for each cohort – all of which used in a genome-wide association with host age. Age-dependent colonization had some heritability, though this varied between cohorts (h^2^ = 0.10, 0.00 – 0.69 95% CI in the first; h^2^ = 0.46, 0.33 – 0.60 95% CI in the second cohort). We found that serotypes and genetic background (strain) explained most of the heritability in each cohort (h^2^_serotype_ = 0.06 and h^2^_GPSC_ = 0.04 in the first; h^2^ _serotype_ = 0.20 and h^2^ _GPSC_ = 0.23 in the second cohort). We found one candidate association (p = 1.2×10^−9^) upstream of an accessory Sec-dependent serine-rich glycoprotein adhesin. Overall, association with age was highly cohort and strain dependent, supporting proposals for a future vaccination strategy which is primarily targeted using serotypes rather than proteins, and is tailored towards specific pathogen populations.

## Introduction

*Streptococcus pneumoniae* is a common commensal of the human upper respiratory tract and nasopharynx, but can also cause invasive diseases such as pneumonia, sepsis or meningitis.(1) Invasive pneumococcal disease (IPD) has a high mortality, and the overall mortality rate from IPD is higher in extreme age ranges, such as infants and the elderly.(2,3) In the Netherlands, pneumococcal carriage rates are higher in children than in adults, with a prevalence of up to 80% at 2 years of age.(4)

Host age is known to affect carriage prevalence and carriage duration of different serotypes(5,6), which is suggested to be driven by differences in immunity.(7) Studies in mice and humans showed evidence for age-dependent host-pathogen interactions involving interleukin (IL)-1 response in reaction to the pore-forming pneumoplysin (*ply*) toxin.(8) IgA secretion is important in clearing *S. pneumoniae* from host upper respiratory tract mucosa and this secretion more effective in previously exposed individuals, the adults.(9) Bacterial genetics has shown to explain over 60% of the variability in carriage duration, and specifically that presence of a bacteriophage inserted in a mediator of genomic competence was associated with a decreased carriage duration.(10)

Pneumococci are highly genetically variable, displaying over 100 diverse capsular serotypes(11), which are a major antigen and the strongest single predictor of carriage prevalence.(12) Pneumococcal conjugate vaccines, targeting up to thirteen capsule serotypes with high burden of invasive disease, cause decreased the rate of nasopharyngeal carriage and invasive disease.(13,14) Besides a direct effect of vaccination with an pneumococcal conjugate vaccine (PCV) on the disease burden in the target population, i.e. young children, it also reduces the disease burden caused by pneumococci with vaccine serotypes in the population not eligible for vaccination through indirect protection from colonization – reducing carriage rates in children reduces overall transmission of the most invasive serotypes.(12,15,16) However, the introduction of PCV has resulted in the replacement of serotypes not covered by the vaccine(17,18), which in some countries reaches levels of invasive disease return towards pre-vaccine levels.(19,20)

As not all serotypes can be included in a conjugate vaccine, three perspectives will lead to improved pneumococcal vaccination have been proposed: whole-cell vaccines(21,22), protein vaccines(23), or changing components in the conjugate vaccine in response to the circulating population.(24) Whole-cell vaccination trials are ongoing, but efficacy remains unproven in human populations.(25) Protein vaccines contain antigens which illicit a strong mucosal immune response, with their targets chosen to be common or conserved in the target population, and ideally reducing onward transmission.(26) In their current form, protein vaccines are not thought to be effective on their own, but if administered with serotype conjugates they may help to reduce serotype replacement. Detailed modelling of the dynamics of pneumococcal population genetics has shown that targeting these vaccines towards serotypes prevalent in specific populations would likely be a superior strategy. This work further shows that providing age-specific vaccine design, using complementary adult-administered vaccines (CAVs) is predicted to have the greatest effect on total IPD burden.(24)

For a future pneumococcal vaccination strategy based on age, we should understand the differences between infant and adult carriage. Differences between host niches have been found, some with a potential effect on onward transmission.(27–29) Treating age as a niche in a systematic analysis of genetic variation would therefore be a powerful way to discover useful vaccine targets. Here, we aim to determine how pathogen variation affects colonization. The identification of age-associated genetic variation, could provide further targets for protein vaccination, whereas ruling these out could provide confidence in age-targeted vaccine formulations based on serotype differences alone.

We performed a pathogen genome-wide association study on pneumococci isolated from nasopharyngeal swabs of 4320 infants and adults from the Netherlands and Myanmar. To dissect pneumococcal variation associated with age we compare prevalence of pneumococcal strains and serotypes between infants and adults in different settings, calculate the contribution of pathogen genetic variation towards predilection for host age, and search for genetic regions associated with host age.

## Methods

### Cohort collection

The Dutch cohort consists of carriage samples from individuals obtained from three prospective carriage surveillance studies.(30–32) In these studies, carriage was assessed by conventional culture of nasopharyngeal or oropharyngeal swabs of vaccinated children (11 and 24 months of age) and their parents in 2009, in 2010/2011, in 2012 and 2013.(30) All children were vaccinated with PCV-7 or PHiD-CV10 according to the Dutch national immunization program at 2, 3, 4 and 11 months of age. Vaccination status of the parents was unknown. Exclusion criteria are described elsewhere.(30,31) Nasopharyngeal swabs were collected from all individuals and oropharyngeal swabs were collected from all adult subjects by trained study personnel using flexible, sterile swabs according to the standard procedures described by the World Health Organization.(33) After sampling, swabs were immediately placed in liquid Amies transport medium and transported to the microbiology laboratory at room temperature and cultured within 12 hours. Pneumococcal isolates were identified using conventional methods, as described previously.(34) The Maela cohort consists of samples from people from a camp for displaced persons on the Thailand-Myanmar border, where monthly nasopharyngeal sampling was performed in unvaccinated children (0 to 24 months old) and their mothers. Procedures for collecting samples and generating whole genome sequences have been previously described.(6,35)

### Informed consent

Written informed consent was obtained from both parents of each child participant and from all adult participants. Approval for the 2009 and 2012/2013 studies in children and their parents (NL24116 and NL40288/NTR3613) were received from the National Ethics Committee in the Netherlands (CCMO and METC Noord-Holland). For the 2010/2011 study, a National Ethics Committee in The Netherlands (STEG-METC, Almere) waived the requirement for EC approval. Informed consent for the Maela cohort was described elsewhere.(6) Studies were conducted in accordance with the European Statements for Good Clinical Practice and the Declaration of Helsinki of the World Medical Association.

### Host age distribution in sequenced carriage cohorts

In the Dutch cohort, children had a median age of 23 months (interquartile range (IQR) 10 – 24 months). Adults had a median age of 35 (IQR 32 – 38) years. In the Maela cohort, the median age of children was 13 months (IQR 6 – 19 months), and for mothers (women of childbearing age) the exact age was unknown (Supplementary Figure S1).(6,36) In the Dutch cohort, all children were vaccinated with PCV-7 or PHiD-CV10. None of the members of the Maela cohort had received PCV.

### DNA extraction and whole genome sequencing

For the Dutch cohort, DNA extraction was performed with the Gentra Puregene Isolation Kit (Qiagen), and quality control procedures were performed to determine yield and purity. Sequencing was performed using multiplexed libraries on the Illumina HiSeq platform to produce paired end reads of 100 nucleotides in length (Illumina, San Diego, CA, USA). Quality control involved analysis of contamination with Kraken (version 1.1.1)(37), number and length of contigs, GC content and N50 parameter. Sequences for which one or more of these quality control parameters deviated by more than 3 standard deviations from the mean were excluded. Sequences were assembled using a standard assembly pipeline.(38) Assembly statistics can be found in the Supplementary Table S1. Genome sequences were annotated with PROKKA, version 1.11.(39) For the Maela cohort, DNA extraction, quality control and whole genome sequencing have been described elsewhere.(40) Serotypes were determined from the whole-genome sequence by in-house scripts.(41) Sequence clusters (strains) were defined as Global Pneumococcal Sequence Clusters (GPSC) using PopPUNK (version 2.2.0), using a previously published reference database.(42,43) For 114 and 401 sequences in the Dutch and Maela cohorts respectively, the GPSC couldn’t be inferred due to low sequence quality.

### Sequencing characteristics and quality control

A total of 1361 bacterial isolates were sequenced as part of the Dutch cohort. During quality control, 32 sequences were excluded. Of these, 8 belonged to a different pathogen species, 9 had contamination, 14 were excluded based on number of contigs or genome length and 1 sequence failed annotation. For 47 sequences, host age was missing. The association analyses were performed on 1282 sequences in the Dutch cohort. Of these, 1052 were isolated from children and 230 from adults. There were 3085 sequences available from the Maela cohort. Quality control for this cohort was described previously.(40) There were 2503 sequences isolated from children and 582 from adults. For the determination of the frequency and odds ratio of serotype and GPSCs in children and adults, only the first isolate from each carriage episode for each child was included in the analysis. This resulted in 964 serotypes and 799 GPSCs (165 missing) in children, and 582 serotypes and 508 GPSCs (74 missing) in adults. For adults Chi-squared tests to calculate the p-value for association between serotype and strain with age were performed in R (version 4.0.0).

### Data availability

Fastq sequences of bacterial isolates from the Dutch cohort were deposited in the European Nucleotide Archive (ENA, study and accession numbers in Supplementary Table S2). Sequences of bacterial isolates in the Maela cohort are available at ENA under study numbers ERP000435, ERP000483, ERP000485, ERP000487, ERP000598 and ERP000599 (Supplementary Table S3).

### Phylogenetic tree

A core genome for sequences from both cohorts together was generated with Roary (version 3.5.0, default parameters), using a 95% sequence identity threshold.(44) A maximum likelihood phylogeny of single-nucleotide polymorphisms (SNPs) in the core genome of all sequenced isolates from both cohorts together was produced with iqtree (version 1.6.5, including fast stochastic tree search algorithm, GTR+I+G) assuming a general time reversible model of nucleotide substitution with a discrete γ-distributed rate heterogeneity and the allowance of invariable sites.(45)

### Heritability analysis

Based on the kinship matrix and phenotypes, a heritability estimate was performed in limix (version 3.0.4 with default parameters) for both cohorts separately.(46) A confidence interval around the heritability estimate was determined with Accurate LMM-based heritability Bootstrap confidence Intervals (ALBI) based on the eigenvalue decomposed distances in the kinship matrix and the heritability estimate with the gglim package (version 0.0.1) in R (version 4.0.0).(47) To estimate the proportion of heritability attributable to serotype or strain alone, we calculated the heritability based on a kinship matrix treating serotypes or strains as genetic variants.(10,48) This analysis was performed in Pyseer (version 1.1.1) using the linear mixed model.(48)

### Determining bacterial genetic variation – unitigs, SNPs and COGs

Using the whole-genome sequence reads from both cohorts, we called SNPs, small insertions and deletions and SNPs clustered as rare variants (deleterious variants at an allele frequency < 0.01) based on the *S. pneumoniae* D39V reference (CP027540) sequence using the Snippy pipeline (version 4.4.0, default parameters). We determined non-redundant sequence elements (unitigs) from assembled sequences in the Dutch cohort by counting nodes on compacted De Bruijn graphs with Unitig-counter (version 1.0.5, default minimum k-mer length of 31).(49) These unitigs were called in an indexed set of sequences from the Maela cohort with Unitig-caller (version 1.0.0, default parameters).(50) This gave us the distribution of sequences from both cohorts with consistent k-mer definitions, making it possible to run predictive models across cohorts. The same Roary run as was used to generate the core-genome alignment was used to extract accessory clusters of orthologous genes (COGs).(44)

There were 966794 unitigs counted from combined sequences in the Dutch cohort. Of these, 303901 passed a minor allele frequency (MAF) of 0.05 filter and had association testing performed. The 9966794 unitigs from the Dutch cohort were called in sequences from the Maela cohort, to obtain 726040 unitigs. Association testing in this group was done for 323112 unitigs which were present at MAF 0.05 or more. Meta-analysis was performed on 251733 overlapping unitigs. There were 313143 SNPs called from sequences in the Dutch cohort, of these 43556 passed MAF filtering. For the Maela cohort, 382230 SNPs were called and 53553 passed the MAF filter. For meta-analysis, 20118 SNPs had overlapping positions and were included. There were 1997 rare variants called in the Dutch cohort, which were burdened in 538 genes. For the Maela cohort, these numbers were 1997 and 423. Together, 186 genes were included in the meta-analysis. Lastly, 2348 COGs were analyzed in the Dutch cohort and 4678 in the Maela cohort. In the meta-analysis there were 627 overlapping COGs.

### Genome-wide association study

The association analysis for SNPs, unitigs, rare variants and COGs was run as a linear mixed model in Pyseer (version 1.1.1), with a minimum minor allele frequency of 0.05.(48) To correct for population structure, the model included a kinship matrix as covariates, which was calculated from the midpoint rooted phylogenetic tree. An association analysis not corrected for population structure was run with unitigs as sequence elements using a simple fixed effects model in Pyseer. Rare variants were clustered in their corresponding gene and analyzed in a burden test. Meta-analysis was performed on summary statistics from the Pyseer results files with METAL (version released on August 28 2018, default parameters) for each variant.(51) A threshold for association of the phenotype with meta-analyzed variants was determined using a Bonferroni correction with alpha < 0.05 and the number of independent tests in the Dutch cohort, giving: p < 1.0×10^−7^ for unitigs, p < 1.0×10^−6^ for SNPs, p < 2.0×10^−5^ for COGs and p < 1.0×10^−4^ for rare variants. Unitigs were mapped to the *S. pneumoniae* D39V reference genome with bowtie-2 (version 2.2.3, with equal quality values and length of seed substrings 7 nucleotides). In accordance with the study populations in both cohorts, the phenotype was dichotomized as host age 0 to 24 months versus adult age (Supplementary Figure S1). Manhattan plots were generated in R, version 3.5.1, with the package ggplot2 (version 3.1.0). Presence or absence of pilus genes was detected by nucleotide BLAST (version 2.6.0, default parameters) analysis. Pilus gene presence association to carriage age was calculated with a likelihood-ratio test in Pyseer (version 1.1.1), corrected for population structure by including a kinship matrix as covariates.

The prediction analysis used the elastic-net mode of Pyseer. This fitted an elastic net model with a default mixing parameter (0.0069 L1/L2) to the unitigs counted in each cohort, using the strains from PopPUNK as folds to try and reduce overfitting.(50) ROC curves for each cohort were drawn using the linear link output, with the R package pROC (version 1.16.2) using smoothing. To test inter-cohort prediction, the called unitigs from the other cohorts were used as predictors with the model from the opposing cohort.

## Results

We first analyzed the observed distribution of serotypes and strains in each of the two cohorts, to assess overall trends of differences between adults and children, and look at the genetic heterogeneity between the two cohorts. Although our cohorts were broadly matched in the primary phenotype, age, large differences between the pathogen population are expected due to different geographies, social backgrounds, and only children in one cohort being vaccinated.

### Serotypes and strains are variable between age groups, and between cohorts

The Dutch cohort was made up of 1329 *S. pneumoniae* isolates comprising 41 unique serotypes (Figure 1, Supplementary Table S4). Of these isolates, 689 (52%) comprised 7 serotypes: 19A (225; 17%), 11A (111; 8%), 6C (97; 7%), 23B (84; 6%), 10A (67; 5%), 16F (54; 4%) and serotype 21 (51; 4%). In this cohort of which the children were vaccinated, a minority of isolates belonged to one of the vaccine serotypes, namely serotype 1 (6; <1%), serotype 4 (1; <1%), serotype 5 (1; <1%), serotype 6B (26; 2%), serotype 7F (11; 1%), serotype 9V (4; <1%), serotype 14 (4; <1%), serotype 18C (5; <1%), serotype 19F (31; 2%) and serotype 23F (12; 1%). The 3085 pneumococcal isolates of the Maela cohort comprised 64 unique serotype groups (Supplementary Table S5). Of these isolates, 1631 (53%) comprised five serotypes: Non-typable (511; 17%), 19F (402, 13%), 23F (307, 10%), 6B (236; 8%) and serotype 14 (175; 6%). In the Dutch cohort, there were 59 unique sequence clusters of which the four largest sequence clusters were GPSC 4 (171; 13%), GPSC 3 (156; 12%), GPSC 7 (131; 10%) and GPSC 11 (119; 9%) (Supplementary Table S6). The. There were 127 unique sequence clusters found in the Maela cohort (Supplementary Table S7). The four largest sequence clusters were GPSC 1 (352; 13%), GPSC 28 (190; 7%), GPSC 20 (168; 6%) and GPSC 42 (123; 5%).

**Figure 1.**
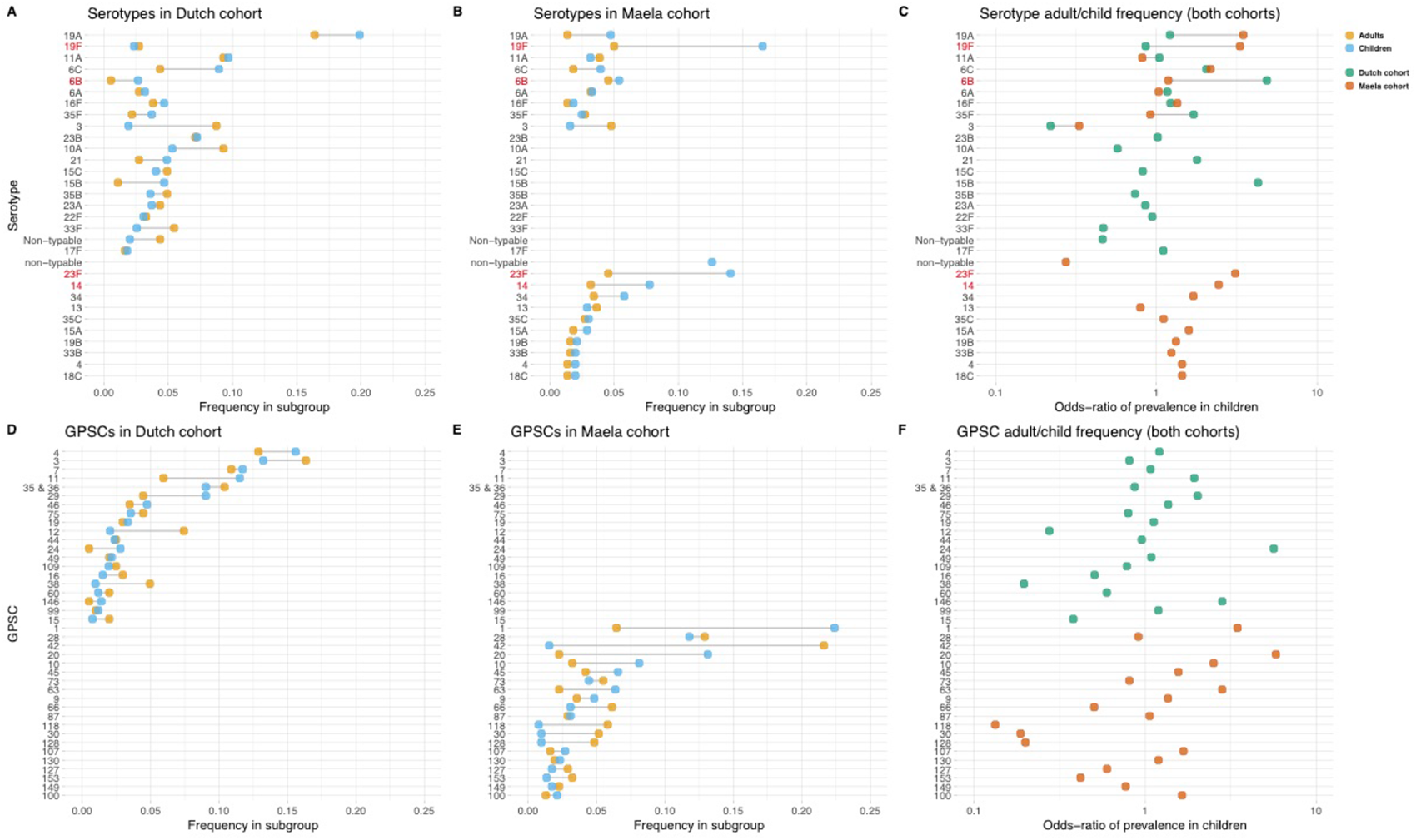
Serotype and strain (GPSC) distribution by age, and between cohorts. Blue dots represent frequency of serotype and strain in child carriage, yellow dots represent frequency in adult carriage. Red and green dots show odds-ratio of prevalence in children in the Dutch and Maela cohorts respectively, on a log scale for serotype. Lines show differences. Top row: dominant serotypes, ordered by presence in cohort, and internally by overall frequency. Vaccine serotypes shown in red. A: Serotype frequency in the Dutch cohort. B: Serotype frequency in the Maela cohort. C: Comparison of adult/child log odds in each cohort for serotype. Second row: dominants strains (GPSCs), ordered by presence in cohort, and internally by overall frequency. D: Strain frequency in Dutch cohort. E: Strain frequency in Maela cohort. F: Comparison of adult/child log odds in each cohort for strain.

Some serotypes exhibited a large difference in colonization frequency between the two age groups. In the Dutch cohort, serotype 6C was overrepresented in children relative to adults (chi-squared test, p = 0.02, not corrected for multiple testing), while in the Maela cohort, serotype groups overrepresented in children were serotype 19F (chi-squared test, p = 3.1×10^−9^), serotype 23F (chi-squared test, p = 1.8×10^−7^), serotype 14 (chi-squared test, p = 1.3×10^−3^); while non-typeable serogroup was overrepresented in adults (chi-squared test, p < 1.0×10^−15^) (Table 1). None of the 20 largest groups of sequence clusters overlapped between the cohorts. In the Dutch cohort only GPSC 11 was significantly associated with carriage in children (chi-squared test, p = 0.03, not corrected for multiple testing). In the Maela cohort, sequence clusters overrepresented in children were GPSC 1 (chi-squared test, p = 1.8×10^−9^) and GPSC 20 (chi-squared test, p = 1.2×10^−7^); while GPSC 42 (chi-squared test, p < 1.0×10^−15^), GPSC 118 (chi-squared test, p = 7.9×10^−5^), GPSC 30 (chi-squared test, p = 9.3×10^−4^) and GPSC 128 (chi-squared test, p = 1.9×10^−3^) were overrepresented in adults (Table 2).

**Table 1.**
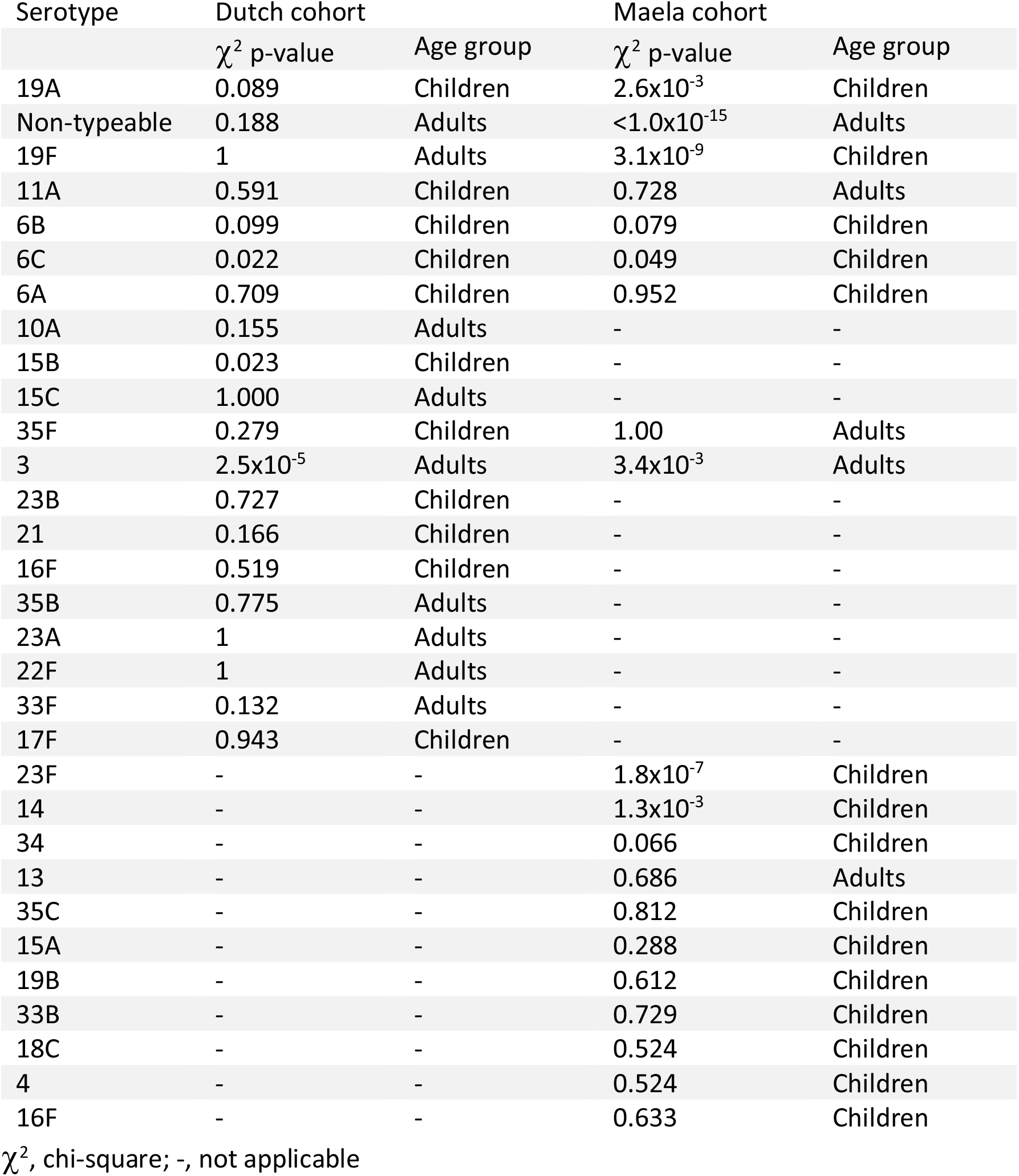
Chi-squared values for serotypes in the Dutch and Maela cohorts and the age group that the serotype is affiliated with

**Table 2.** Chi-squared values for strains in the Dutch and Maela cohorts and the age group that the strain is affiliated with

| GPSC | Dutch cohort |  | Maela cohort |  |
| --- | --- | --- | --- | --- |
| | $\chi^2$ p-value | Age group | $\chi^2$ p-value | Age group |
| 60 | 0.568 | Adults | - | - |
| 4 | 0.298 | Children | - | - |
| 3 | 0.392 | Adults | - | - |
| 7 | 0.858 | Children | - | - |
| 11 | 0.03 | Children | - | - |
| 35 & 36 | 0.617 | Adults | - | - |
| 29 | 0.049 | Children | - | - |
| 46 | 0.563 | Children | - | - |
| 75 | 0.666 | Adults | - | - |
| 19 | 0.978 | Children | - | - |
| 12 | $1.2 \times 10^{-4}$ | Adults | - | - |
| 44 | 1 | Adults | - | - |
| 24 | 0.094 | Children | - | - |
| 49 | 1 | Children | - | - |
| 109 | 0.817 | Adults | - | - |
| 16 | 0.249 | Adults | - | - |
| 38 | $2.1 \times 10^{-4}$ | Adults | - | - |
| 146 | 0.489 | Children | - | - |
| 99 | 1 | Children | - | - |
| 15 | 0.22 | Adults | - | - |
| 1 | - | - | $1.8 \times 10^{-9}$ | Children |
| 28 | - | - | 0.959 | Adults |
| 20 | - | - | $1.3 \times 10^{-7}$ | Children |
| 42 | - | - | $< 1.0 \times 10^{-15}$ | Adults |
| 10 | - | - | 0.005 | Children |
| 45 | - | - | 0.1462 | Children |
| 63 | - | - | 0.008 | Children |
| 73 | - | - | 0.754 | Adults |
| 9 | - | - | 0.388 | Children |
| 66 | - | - | 0.086 | Adults |
| 87 | - | - | 0.928 | Children |
| 118 | - | - | $7.9 \times 10^{-5}$ | Adults |
| 30 | - | - | $9.3 \times 10^{-4}$ | Adults |
| 128 | - | - | $1.9 \times 10^{-3}$ | Adults |
| 107 | - | - | 0.372 | Children |
| 127 | - | - | 0.464 | Adults |
| 130 | - | - | 0.809 | Children |
| 153 | - | - | 0.147 | Adults |
| 149 | - | - | 0.885 | Adults |
| 100 | - | - | 0.479 | Children |

$\chi^2$ , chi-square; -, not applicable

A phylogenetic tree of pooled sequences from both cohorts, with serotype, sequence cluster, age group and cohort for each sequence, revealed clonal discrimination between cohorts (Figure 2). Combined with the effects shown in Figure 1, this highlighted a key feature of our analysis of these datasets, which was the genetic heterogeneity between the two cohorts. Individually, each dataset clearly has strains and serotypes with strong signals of host age differences, but the overall makeup of each dataset is very different (twelve common serotypes are shared, but only a single common GPSC), and where there are shared serotypes many have different effect directions between the two cohorts.

**Figure 2.**
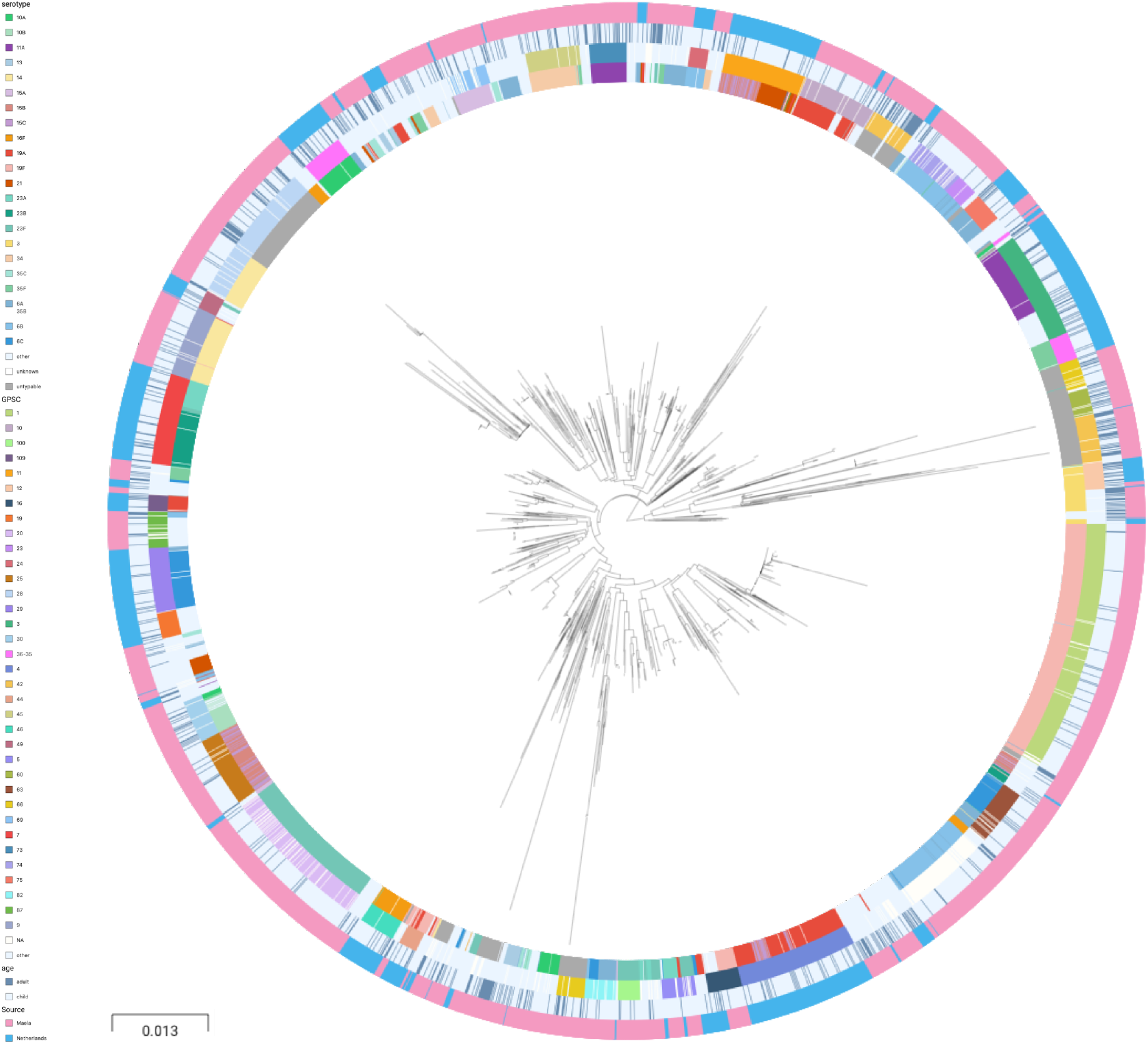
Phylogenetic tree of carriage samples from both cohorts. The rings show metadata for the samples. Depicted from inside to outside, these are serotype, sequence cluster (GPSC), age and source (Maela, Netherlands). Scale bar: 0.013 substitutions per site. An interactive version is available at https://microreact.org/project/f2MdBLZhSyU9eF8MBobHhA/e2a5ebd7 (project link https://microreact.org/project/f2MdBLZhSyU9eF8MBobHhA).

### Host age is heritable and mostly explained by strain and serotype

To quantify the amount of variability in carriage age explained by variability in the genome, we calculated a heritability estimate (h^2^) for each cohort. For isolates in the Dutch cohort, we did not find strong evidence that genetic variability in bacteria was related to variance in host age (h^2^ = 0.10, 0.00 – 0.69 95% CI). In the Maela cohort, we found significant evidence that affinity with host age was heritable (h^2^= 0.46 0.33 – 0.60 95% CI) and thus genetic variation in this cohort explained variation in carriage age to a greater degree. In both cohorts pan-genomic variation could be used to predict host age to some degree of accuracy (area under the ROC curve 0.82 [Dutch cohort]; 0.91 [Maela cohort]), suggestive of some level of heritability and association of host age with strain (Figure 3). Prediction between cohorts using a simple linear model failed, as the genetic variants chosen as predictors were not found in the other cohort – again highlighting the high level of genetic heterogeneity between cohorts.

**Figure 3.**
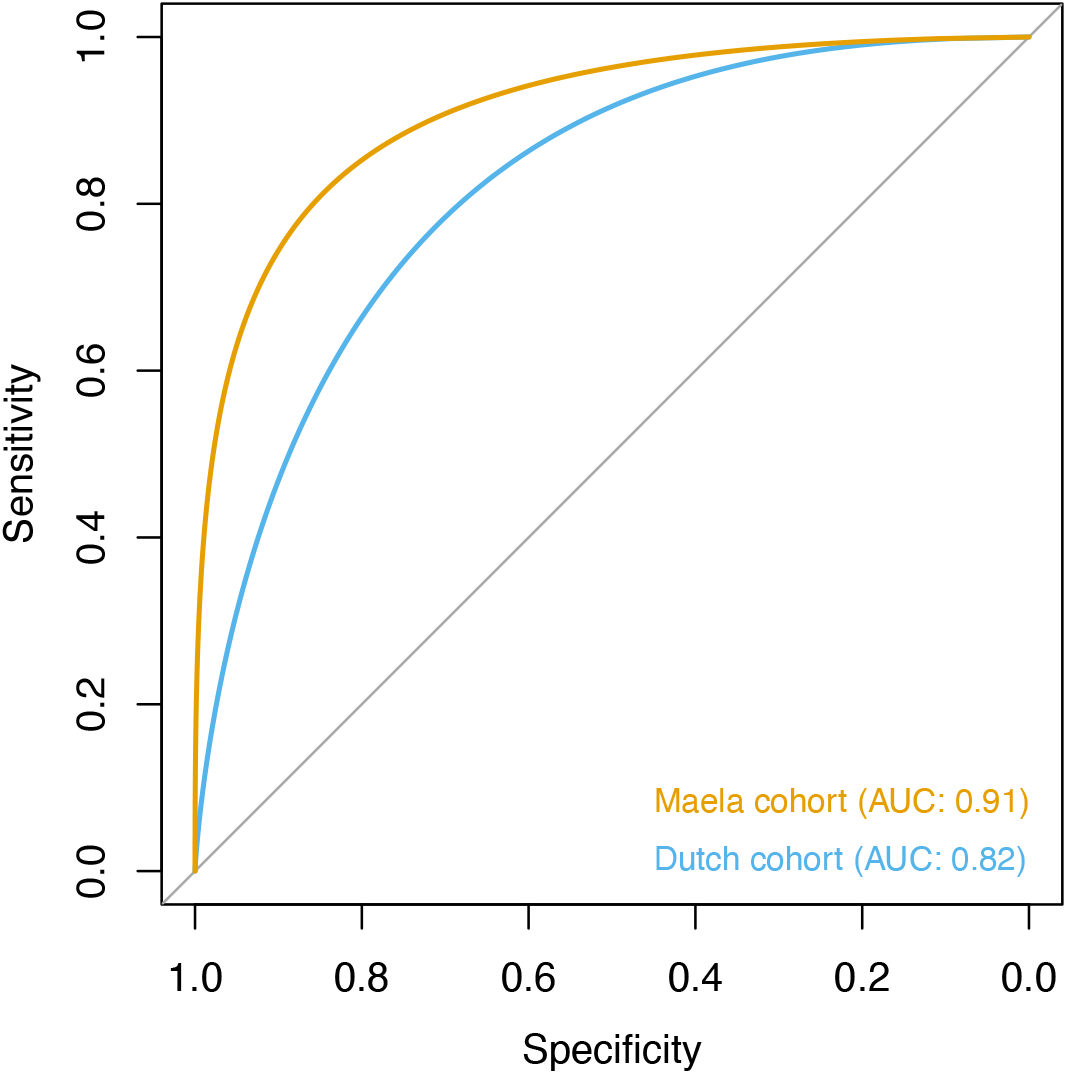
Prediction of host age from pan-genomic variation in each cohort. The smoothed ROC curve based on a linear predictor (elastic net fitted to unitigs, with strains used as folds for cross-validation) is shown. Area under the curve (AUC) is 0.5 for no predictive ability and 1 for perfect prediction.

To further investigate the association of serotype and sequence cluster to carriage age, we determined the proportion of variation in carriage age explained by serotype and sequence cluster alone. Here, we estimated *h*^2^_serotype_ = 0.06 and *h*^2^_GPSC_ = 0.04 for the Dutch cohort and *h*^2^ _rotype_ = 0.20 and *h*^2^_GPSC_= 0.23 for the Maela cohort, confirming the larger contribution of serotype and sequence cluster to carriage age in sequences from the Maela cohort. We also performed a genome-wide association analysis, but without controlling for population structure. This reveals genetic variants specific to serotype as determinants for carriage age (p-values < 5.0×10^−8^) in both cohorts (Supplementary Table S8 and Supplementary Table S9). Among the genetic variants with the lowest p-values were variants in capsule locus genes (Cps) in both cohorts. This further supports a role of strain and serotype in association with host age, but does not distinguish between the two.

### Genome-wide association analysis does not find genetic variants independent of strain

Following these observations that serotype and strain do not explain the full heritability, we performed a pathogen genome-wide association analysis to investigate whether we can detect genetic variants irrespective of the genetic background that are associated with carriage in children or adults. Though the cohorts have little genetic overlap in terms of genetic background, we would be well-powered to detect genetic variation independent of background (‘locus’ associations).(52,53) In the Dutch cohort, none of the unitigs, SNPs, COGs or rare variants surpassed the threshold for multiple testing correction (Supplementary Figure 1). The burden (sum) of rare variants in a gene for tryptophan synthase, *trpB*, approach the multiple testing threshold, but was not significant. In the Maela cohort unitigs in the *ugpA* gene surpassed the threshold for statistical significance (Supplementary Figure 2). After meta-analysis, there were two hits which surpass the threshold for statistical significance (Figure 4).

**Figure 4.**
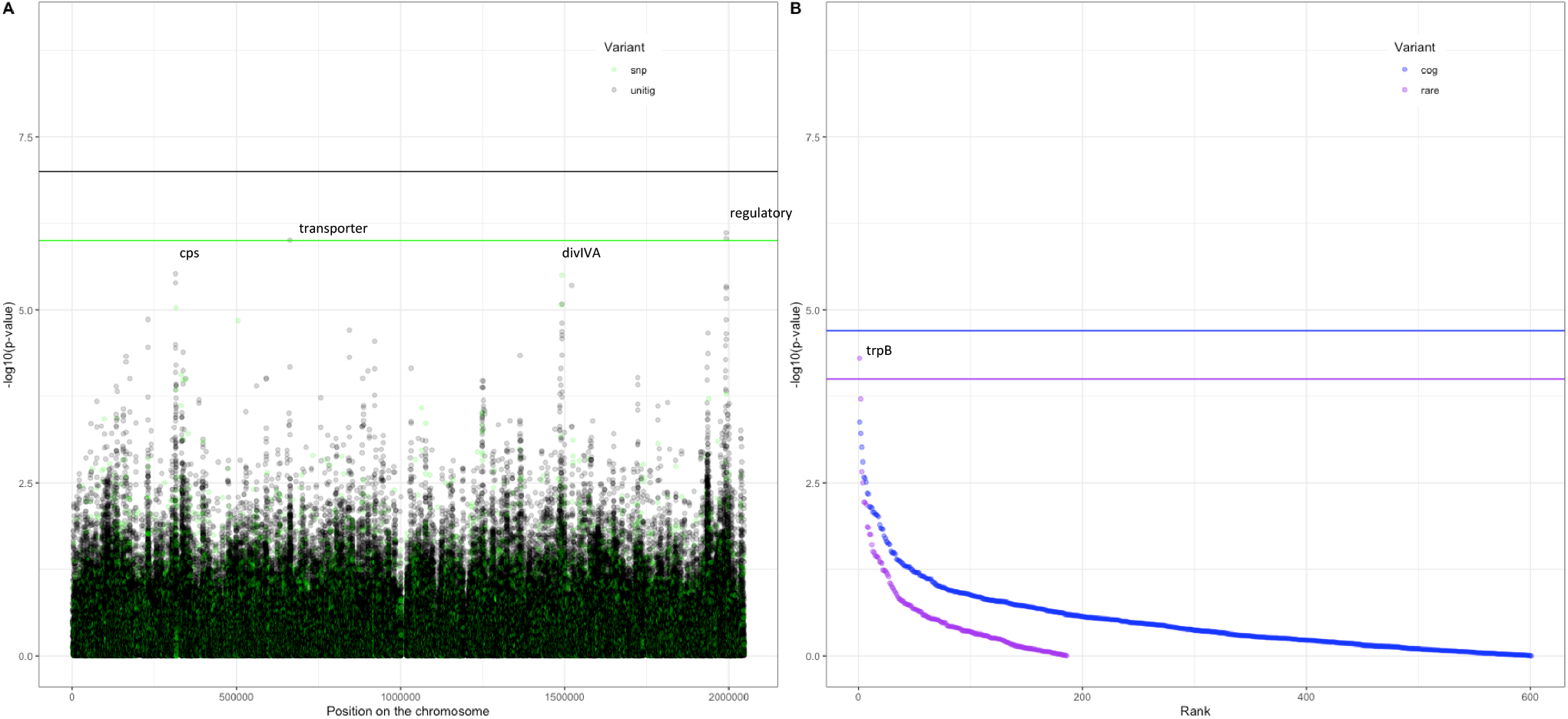
Association of variants after meta-analysis with carriage age 0 – 24 months, A. minus log transformed p-value on the y-axis and position of unitig and snp variants on the *S. pneumoniae* genome on the x-axis (Manhattan plot); B. minus log transformed p-value on the y-axis and sorted lowest to highest p-value for rare variant burden in genes (purple) and clusters of orthologous genes (COGs, blue) on the x-axis.

The first, a nucleotide sequence marked by multiple unitigs of which the lowest has a p-value of 1.2×10^−9^ (Supplementary Table S10). This sequence does not map to the *S. pneumoniae* D39V reference sequence.(54) For this reason, it is not visualized on the Manhattan plot, for which unitigs were mapped to the *S. pneumoniae* D39V reference sequence. Upon inspection of the individual sequences these unitigs are called from, we find them to map in the intergenic region between open reading frames encoding the accessory Sec-dependent serine-rich glycoprotein adhesin and a MarR like regulator, respectively. This region contains sequences resembling transposable elements and an open reading frame encoding a transposase. The unitigs map upstream of the start codon of the accessory Sec-dependent serine-rich glycoprotein adhesin. The sequence is present in 169 out of 1282 (13%) sequences in the Dutch cohort and in 241 out of 3085 (8%) in the Maela cohort. The sequence is present in isolates dispersed over the phylogenetic tree, and associated with carriage in children (Supplementary Figure S4). This protein is involved in adhesion to epithelial cells and biofilm formation.(55–57) Given that this sequence lies just upstream of the start codon, it is plausible that variation of this sequence alters the expression of the Sec-dependent adhesin protein, and therefore affects carriage.

The second hit is a burden of rare variants in a gene for tryptophan synthase, *trpB*, that surpass the threshold for statistical significance at a p-value of 5.0×10^−5^. The variants are 2 frameshift variants of very low frequency. These result in a predicted dysfunctional *trpB* gene in 9 out of 1282 (1%) sequences in the Dutch cohort and in 12 out of 3073 (0.4%) sequences in the Maela cohort. This association of the *trpB* gene is likely to be an artefact of low allele frequency, as we estimate we are only powered to detect variation in at least 5% of isolates.

### Pilus gene presence does not determine carriage age independent of genetic background

Finally, we investigated whether pneumococcal isolates containing a pilus gene preferentially colonize children in the Dutch cohort, as has been previously described in the Maela cohort.(9,58) This study analyzed the Maela cohort and found that 934 out of 2557 (37%) isolates in children versus 95 out of 592 (16%) isolates in adults had pilus genes present. However, this association of pilus gene presence to carriage age was dependent on lineages within the population.(9) In the Dutch cohort, we found no evidence that host age was dependent on pilus gene presence (22 out of 208 (10%) in adults versus 129 out of 1099 (12%) in children). This was the case whether or not the genetic background was adjusted for (p = 0.35, uncorrected for population structure and p = 0.69, corrected for population structure). Based on these findings, we suggest that the previously reported pilus-IgA1 association is not a universal explanation for difference in colonization between hosts of different ages.

## Discussion

The age of the host is known to have an important effect on pneumococcal colonization.(12) Observational studies have demonstrated variation in serotype prevalence and carriage duration between infants and adults. Mechanistic studies in mice and humans have shown examples of differing immune responses depending both on host factors and pathogen factors. Findings from these studies include the observation that capsular polysaccharides (determinants of serotype) inhibit phagocytic clearance in animal models of upper respiratory tract colonization.(59) A pneumolysin-induced IL-1 response determined colonization persistence in an age-dependent manner(8); and pilus expressing strains were found to preferentially colonize children, because of immune exclusion via secretory IgA in non-naïve hosts.(9)

Building upon these observations, we sought to investigate and quantify the contribution of pathogen genetic variation to carriage in infant versus adult hosts, using a top-down approach. Through whole genome sequencing and application of statistical genetic methods to two large *S. pneumoniae* carriage cohorts, we show evidence that bacterial genetic variability influences predilection for host age, though this appears to be highly variable between populations. One important difference between our study cohorts was that children from the Dutch cohort were vaccinated, while children from the Maela cohort were not. While our findings demonstrate that vaccinated versus unvaccinated children were colonized with different bacterial serotypes and different sequence clusters, we observed differences in prevalence beyond just the serotypes included in the vaccine. Another difference between the cohorts was that adults from the Dutch cohort were males and females, while adults from the Maela cohort were female only.

Strain, or genetic background, appears to be the main effect, explaining roughly half of the total heritability in each cohort. We found sequences in one region which map closely to the start codon of the accessory Sec-dependent serine-rich glycoprotein adhesin to be associated with carriage age independent of genetic background, in a meta-analysis of the two cohorts. Our results our suggestive of a polygenic architecture of many variants with low effect sizes, along with larger effects between strains. Three reasons can contribute to this: the proportion of the heritability which is caused by lineage effects; rare locus effects which could not be detected with the current sample size; and by sampling from a cohort with vaccinated children and unvaccinated adults and comparing with a cohort of unvaccinated children and adults, we had lower power due to the reduced overlap within and between cohorts in pan-genome content. Although differences in vaccination status between cohorts is a plausible explanation for our findings, we were unable to rule out other factors, for example a population-specific host effect, or the broad effects of different socio-economic status between these cohorts.

In previous bacterial GWAS studies of antimicrobial resistance (such as a single gene which causes antibiotic resistance), large monogenic effects have typically been found to have high heritabilities close to one, and the GWAS identify the causal variant precisely.(50,52,60) When applied to virulence and carriage duration phenotypes, heritable effects have also been found, but these only explained some of the variation in the phenotype. These appeared to be caused by weaker polygenic effects, not all of which could be detected using the relatively small cohorts available.(10,61) We found similar results for host-age heritability in these two cohorts. Within this genetic architecture, our finding of a sequence just upstream of the start codon of the accessory Sec-dependent serine-rich glycoprotein adhesin being associated with carriage in children, is notable.

We could not distinguish between genetic background or serotype being the primary effect due to their correlation. We did note a difference in effect size of serotype between the two cohorts, which may make it unlikely to be the single largest effect on host age. This difference in cohorts could be explained by strain/GPSC being the main and consistent effect on host age. As strains are different between cohorts and each serotype appears in multiple strains, combining them in different amounts would create different directions of effect for serotype. We did not replicate the association of piliated genomes in infant hosts in our newly sequenced cohort, further demonstrating important differences between populations.

In summary, we found an effect of pneumococcal genetics on carriage in children versus adult hosts, which varies between cohorts, and is likely primarily driven by strain (lineage) effects rather than large population-wide effects of individual genes. An important corollary of our work is on future pneumococcal vaccine optimization efforts. A promising approach for future vaccination strategies is to target the different age groups.(24) Whether these should consist of the dominant disease-causing serotypes overrepresented in carriage by each age group, or whether there are age-specific pathogen proteins that should be included is an open question. Our study suggests that targeting these age groups using serotype makeup alone would be sufficient, and supports previous observational and modelling studies which advise targeting the serotype makeup in the vaccine at specific populations to maximize their effect.

## Acknowledgments

We would like to thank Dr. Nicholas Croucher from Imperial College London for commenting on the manuscript.

