## Supplementary figures and images for "Pneumococcal genetic variability influences age-dependent bacterial carriage"

### Supplemental Figure S1

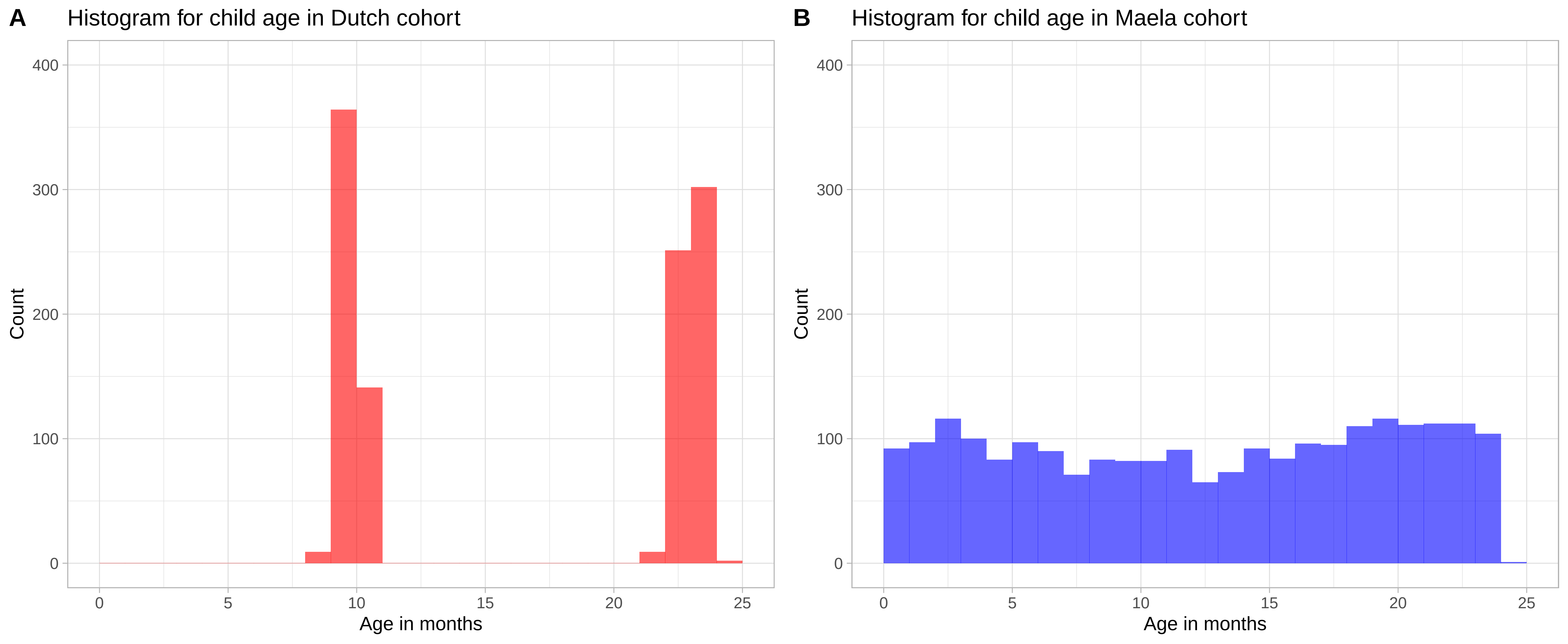

### Supplemental Figure S2

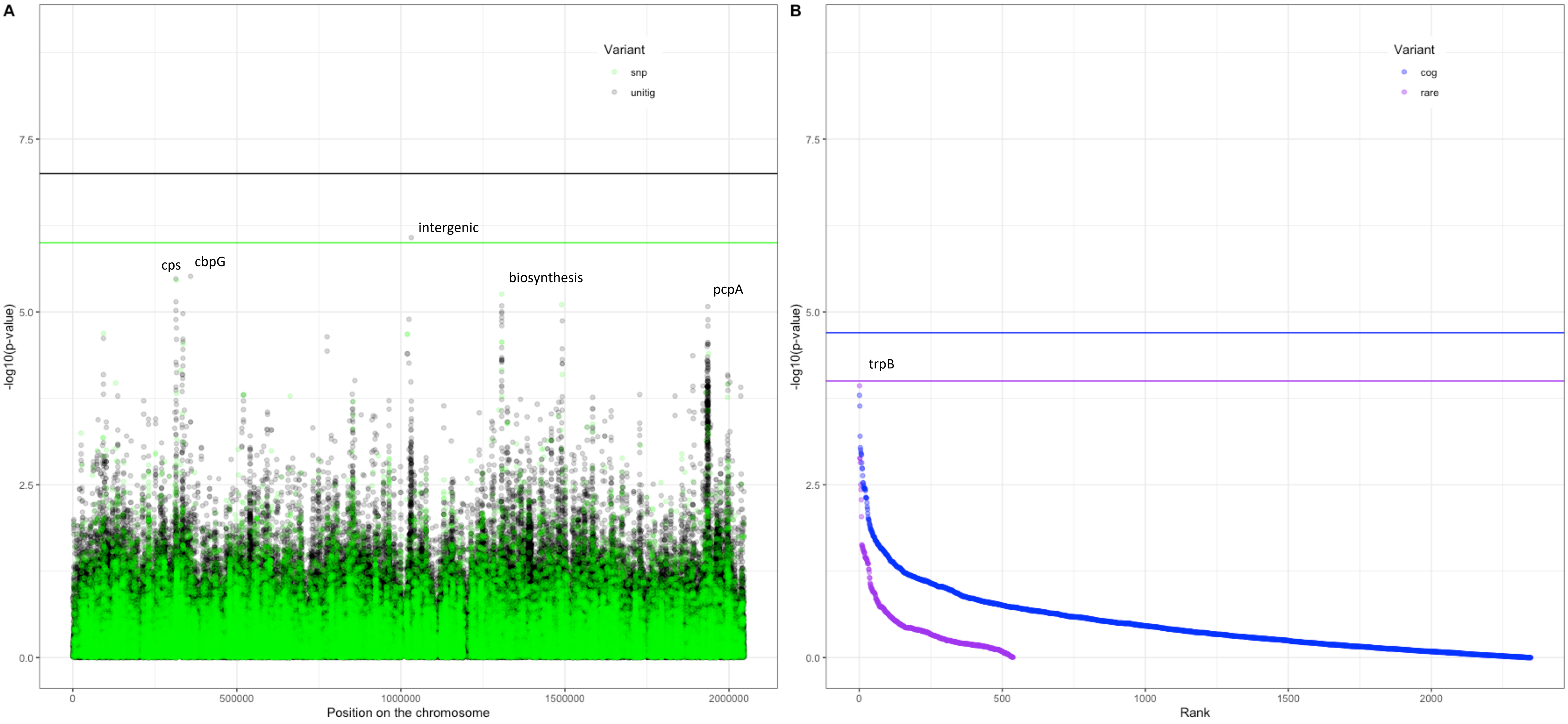

### Supplemental Figure S3

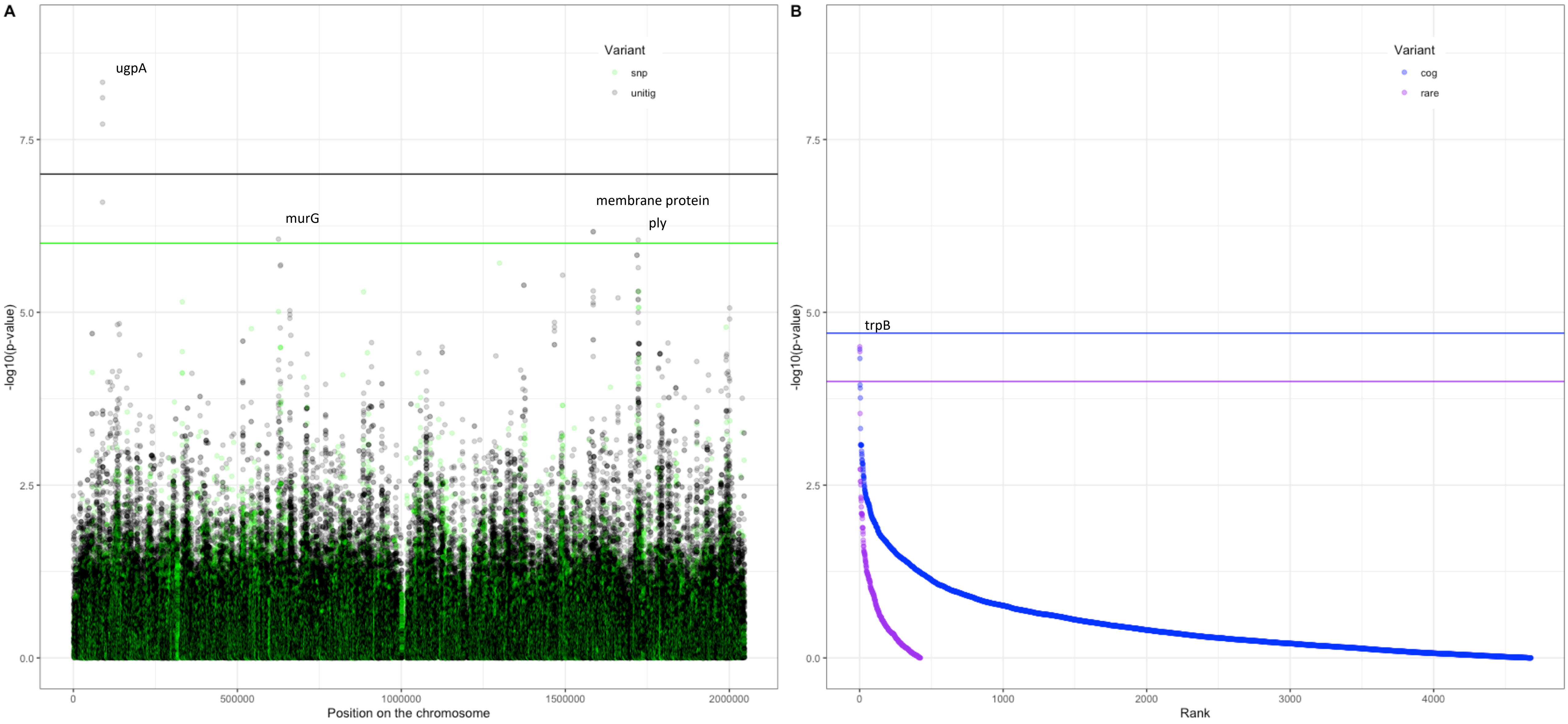

### Supplemental Figure S4

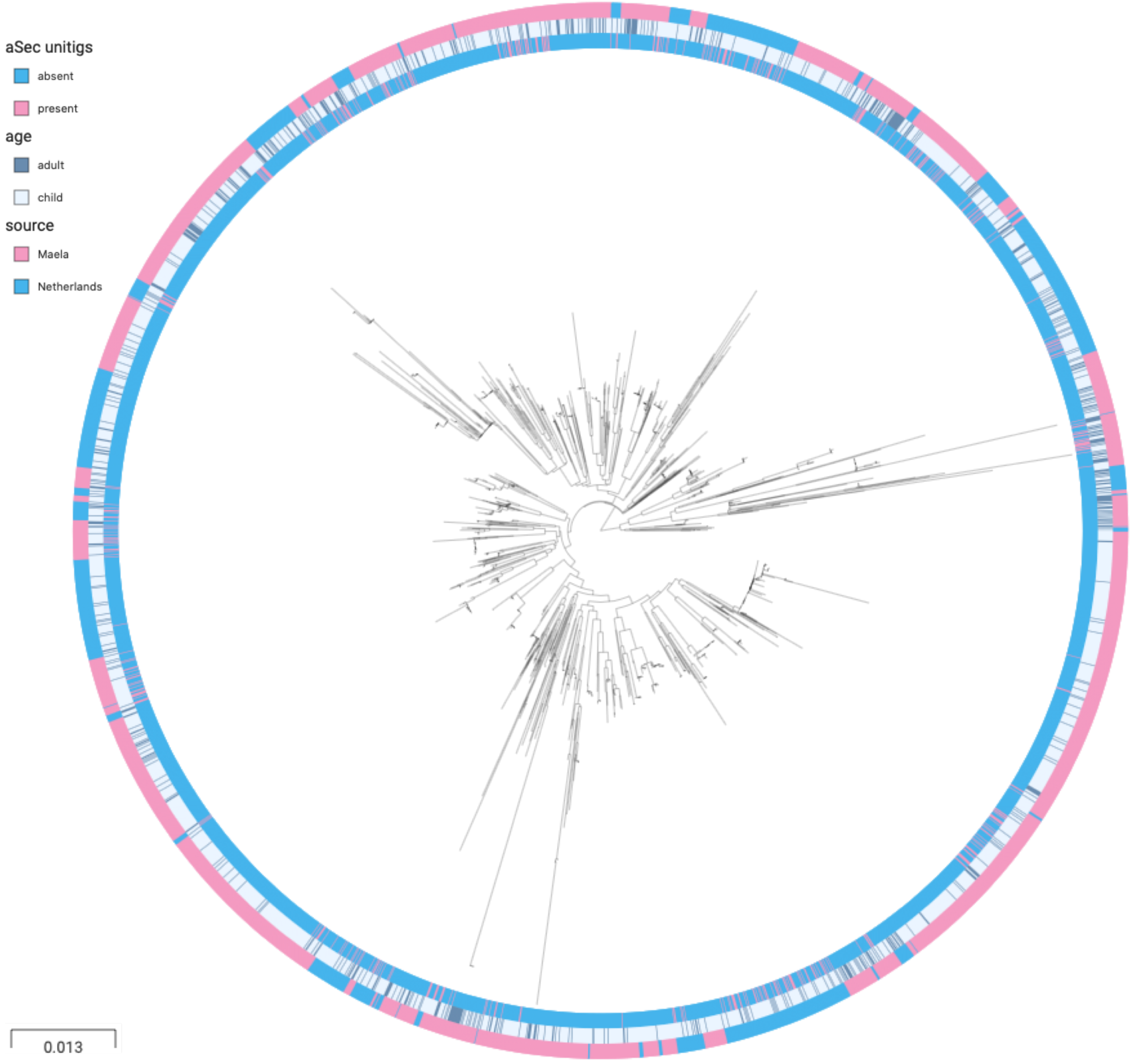
